# Defining the pig microglial transcriptome reveals their core signature, regional heterogeneity, and similarity with humans

**DOI:** 10.1101/2021.08.11.454467

**Authors:** Barbara B Shih, Sarah M Brown, Lucas Lefevre, Neil A Mabbott, Josef Priller, Gerard Thompson, Alistair B Lawrence, Barry W McColl

**Author notes:** These authors contributed equally (Barbara B Shih, Sarah M Brown, Alistair B Lawrence, Barry W McColl).

## Abstract

Microglia play key roles in brain homeostasis as well as responses to neurodegeneration and neuroinflammatory processes caused by physical disease and psychosocial stress. The pig is a physiologically-relevant model species for studying human neurological disorders, many of which are associated with microglial dysfunction. Furthermore, pigs are an important agricultural species, and there is a need to understand how microglial function affects their welfare. As a basis for improved understanding to enhance biomedical and agricultural research, we sought to characterise pig microglial identity at genome-wide scale and conduct inter-species comparisons.

We isolated pig hippocampal tissue and microglia from frontal cortex, hippocampus and cerebellum, as well as alveolar macrophages from the lungs and conducted RNA-sequencing (RNAseq). By comparing the transcriptomic profiles between microglia, macrophages, and hippocampal tissue, we derived a set of 365 highly-enriched genes defining the porcine core microglial signature. We found brain regional heterogeneity based on 215 genes showing significant (adjusted p<0.01) regional variations and that cerebellar microglia were most distinct. We compared normalized gene expression for microglia from human, mice and pigs using microglia signature gene lists derived from each species and demonstrated that a core microglial marker gene signature is conserved across species, but that species-specific expression subsets also exist. Importantly, pig and human microglia shared greater similarity than pig and murine microglia.

Our data provide a valuable resource defining the pig microglial transcriptome signature that highlights pigs as a useful large animal species bridging between rodents and humans in which to study the role of microglia during homeostasis and disease.

**Main Points:**

- Defined a pig microglial transcriptome signature comprising 365 genes.
- Demonstrated regional variance in the pig microglial transcriptome across the brain.
- Revealed greater similarity between pig and human microglia than mouse.

## Introduction

Microglia are resident mononuclear phagocytes of the central nervous system (CNS) parenchyma that are increasingly recognised to play an important role in the development, homeostasis and diseases of the CNS (Li & Barres, 2018).

Microglia are derived from erythro-myeloid progenitors during early embryonic development (Reemst, Noctor, Lucassen, & Hol, 2016) and are not replenished by blood monocytes under normal physiological conditions (Gomez Perdiguero et al., 2015). Microglia sense changes in their environment through a large repertoire of receptors, and mediate responses that promote neuronal and synaptic health, and assist in tissue protection and repair to microbial and sterile injury stressors (S. Hickman, Izzy, Sen, Morsett, & El Khoury, 2018). However, in specific contexts some phenotypes of microglia are thought to contribute to disease processes including in neurodegenerative disease. Indeed, reactive microglia and inflammatory cytokines are commonly observed around lesions in several neurodegenerative disorders, including Alzheimer’s disease (Frautschy et al., 1998), Parkinson’s disease (McGeer, Itagaki, Boyes, & McGeer, 1988), and multiple sclerosis (Kuhlmann et al., 2017). Age-dependent changes in microglia activation and regulation have been reported in rodents (Ogura, Ogawa, & Yoshida, 1994; Perry, Matyszak, & Fearn, 1993), nonhuman primates (Sheffield & Berman, 1998) and humans (Streit & Sparks, 1997), and have been associated with deficits in psychomotor coordination (Richwine et al., 2005) and cognitive function (Jang, Dilger, & Johnson, 2010; Rosczyk, Sparkman, & Johnson, 2008) in mouse models of aging. Microglial reactivity is also associated with a wide range of psychosocial stressors (Calcia et al., 2016; Stein, Vasconcelos, Albrechet-Souza, Cereser, & de Almeida, 2017), and behavioural susceptibility to social stress is driven by microglial induced increases in reactive oxygen species (Lehmann, Weigel, Poffenberger, & Herkenham, 2019).

Microglia have a distinct transcriptome from tissue-resident macrophages in other organs and from the other cell types in the CNS (Butovsky et al., 2014). A number of transcriptomic studies have characterised the gene expression signature for microglia in non-neuropathologic individuals in humans and mice (Bennett et al., 2016; Butovsky et al., 2014; Darmanis et al., 2015; Galatro et al., 2017; Hawrylycz et al., 2012; Sankowski et al., 2019). These microglial gene signatures have been instrumental to understanding the core molecular identity of microglia, their diversity, and as a basis to characterise the spatiotemporal transcriptional changes of microglia in response to aging and disease conditions (Hammond et al., 2019; Patir, Shih, McColl, & Freeman, 2019). Microglial studies have generally been conducted in rodent models or human post-mortem brain tissue. While many cross-species similarities are evident, differences have also been described [19, 22], and may have implications when extrapolating findings across different animal species. Significant differences may also be evident when comparing species such as mice and humans due to substantial disparities in body weight, brain mass and lifespan, which can be partly mitigated in some large animal species.

Pigs are an attractive, physiologically relevant animal model for studying human neurological disorders. In contrast to rodents, pigs have large human-like gyrencephalic brains and human-like grey:white matter ratio. These anatomic features are ideally suited for neuroimaging, cell transplantation and gene therapy studies (Lind et al., 2007; Sauleau, Lapouble, Val-Laillet, & Malbert, 2009; Simchick et al., 2019; Sjöstedt et al., 2020). Recent data from Sjöstedt et al. (2020) suggested that the global gene expression profiles for some brain regions (such as the cerebellum and hypothalamus) in pigs were more similar to those of humans, than those of mice to humans (Sjöstedt et al., 2020). However, the transcriptome-wide signature that specifically defines microglia in the pig brain is not known. As microglia are the principal cellular mediators of innate immunity in the CNS, it is relevant to note previous studies indicating the greater similarity of pigs (than rodents) to humans in certain aspects of innate immune physiology, notably macrophage activation signalling (Fairbairn, Kapetanovic, Sester, & Hume, 2011; Kapetanovic et al., 2012). Transcriptional analyses of pig mononuclear phagocytes have suggested their responses are more similar to human cells than those from mice (Robert et al., 2015). E.g. pigs and other large mammals differ from mice in their ability to induce the expression of genes responsible for arginine metabolism and nitric oxide production (Bush et al., 2020). Furthermore, microglia are thought to be instrumental in mediating responses to non-disease challenges such as social stress (Mondelli, Vernon, Turkheimer, Dazzan, & Pariante, 2017; Salter & Stevens, 2017). With the potential to harness the pig as an intermediate species for translational biomedical research, it is important to establish a normative microglial profile in pigs and instructive to relate this to signatures in other species. In addition, pigs are one of the most economically important and intensively farmed livestock species, and a better molecular definition of pig microglia may aid understanding of conditions that can promote their health, welfare and productivity. Our previous work implicated microglia in the effects of environmental enrichment on neural health including altered microglial gene expression in pigs (Brown, Bush, Summers, Hume, & Lawrence, 2018). The aims of this study were to define the transcriptome identity of pig microglia, and conduct comparative analysis across brain regions, and with mouse and human microglia signatures.

## Methods

### Ethical review

All work was carried out in accordance with the U.K. Animals (Scientific Procedures) Act 1986 under EU Directive 2010/63/EU following ethical approval by SRUC (Scotland’s Rural College) Animal Experiments Committee. All routine animal management procedures were adhered to by trained staff and health issues treated as required. All piglets remaining at the end of the study were returned to commercial stock.

### Animals and general experimental procedures

Sixteen commercial cross-bred female breeding pigs (sows; Large White x Landrace) were artificially inseminated using commercially available pooled semen (Danish Duroc). Piglets were born into either standard commercial housing or pens, allowing greater behavioural freedom (Baxter, Lawrence, & Edwards, 2011). Sows were balanced for parity across both conditions. No tooth resection was performed and males were not castrated. In line with EU Council Directive 2008/120/EC tail docking was not performed. In accordance with the Defra Code of Recommendations for the Welfare of Livestock, temperature within the room was automatically controlled at 22°C and artificial lighting was maintained between the hours of 0800 to 1600, with low level night lighting at other times. At around 21 days of age small amounts of weaning diet (ForFarmers Ultima 2) was introduced to the piglets. At between 24 and 26 days of age, one male piglet (7-8 kg) per litter was selected for tissue collection. The piglets used in this study were part of a wider study involving *in vivo* neuroimaging by MRI under sedation. Piglets were sedated with a combination of ketamine (5 mg/kg), midazolam (0.25 mg/kg) and medetomidine (5 micrograms/kg) injected intra-muscularly (quadriceps). After 3 – 5 min when profound sedation was present, anaesthesia was induced with 2 – 3 % isoflurane delivered by a Hall pattern mask until adequate jaw relaxation allowed laryngoscopy and the topical application of 0.8 – 1.0 mL 2% lidocaine solution. endotracheal intubation was conducted using a 5 mm OD endotracheal tube 90 s later. Anaesthesia was maintained with isoflurane in O_2_ delivered using a Bain breathing system. The lungs were inflated mechanically to produce normocapnia. Core temperature was maintained using hot air blowers. The animal was euthanised humanely under anaesthesia using pentobarbital IV (40 mg/kg) IV.

### Tissue collection

Piglet brains were removed and then cut into two hemispheres. All dissections were performed by a single experienced researcher using http://www.anatomie-amsterdam.nl/sub_sites/pig_brain_atlas for reference, utilising both parasagittal and rostrocaudal views. One hemisphere was dissected into broad anatomical regions for microglial isolation and tissue RNA extraction. Samples for microglial isolation were minced in 1X HBSS (w/o Ca^2+^ and Mg^2+^, 12mM HEPES) and placed on ice for immediate cell isolation. Adjacent samples for tissue level RNA extraction were placed in RNAlater at room temperature for 30 min then snap frozen on dry ice and stored at −20°C until required. The average time from confirmation of death to tissue being stored in RNAlater was approximately 8.5 min. The opposite hemisphere was placed directly into 4% PFA for later histological analysis. After 10 days the solution was changed for Tris Azide and samples maintained at 4°C. Alveolar macrophages were collected in saline solution by post-mortem bronchoalveolar lavage (< 5 min from confirmation of death) and placed on ice for immediate cell isolation (further 1-2 min).

### Preparation of brain cell suspensions

We adapted methods previously described for rodent microglial isolation (Grabert & McColl, 2018). Samples of frontal cortex, hippocampus and cerebellum in 1XHBSS buffer were transferred individually to a glass Dounce homogeniser on ice and cells dissociated by 40 passes of the pestle. Cells were pelleted by centrifugation and resuspended in 35% isotonic Percoll over ice. The sample was overlaid with 1XHBSS without disrupting the Percoll gradient and spun at 4°C for 45 min. The separated myelin layer and supernatant were removed and the cell pellet resuspended in 10 ml HBSS. Cell suspensions were filtered through a 70 μm filter and pellets resuspended in FACS (fluorescence-activated cell sorting) buffer (1X PBS, 25mM HEPES, 0.1% BSA).

### Isolation of microglia and alveolar macrophages

Brain and alveolar lavage cell suspensions, were incubated with 1% Human IgG1 Fc block (R&D systems) for 30 min. Samples were centrifuged, supernatant removed and cells resuspended in a combination of mouse anti-human CD11b:Pacific Blue (Biolegend) and mouse anti-pig CD45:AF647 antibodies (BioRad) for microglial isolation, or mouse anti-pig CD163:FITC (Biorad) and F4/80:AF647 (mouse anti-pig ADGRE1; ROS-4E12-3E6)(Waddell et al., 2018) antibodies for alveolar macrophage isolation for 30 min incubation at room temperature. Samples were centrifuged, supernatant removed and cells resuspended in FACS buffer. Samples were sorted on a FACSAria III cell sorter. Microglia in brain cell suspensions were identified as CD11b^hi^CD45^lo^ and macrophages identified from the alveolar lavage suspensions as F4/80^+^CD163^lo^. Sorted cells were collected directly into TRIzol reagent (Invitrogen) for immediate RNA isolation.

### Microglial and Macrophage RNA extraction

Cell suspensions were transferred into 2 ml lysing matrix D tubes (MP Biomedicals) and homogenised on a FastPrep 24 at 6.5m/s for 50 s. Homogenate was transferred to a 1.5 ml microtube and incubated at room temperature for 5 min to allow complete dissociation of nucleoprotein complexes. An equal volume of chloroform was added to the homogenate, samples placed on a shaker at room temperature for 5 min, and then centrifuged at 12000rmp, 4°C for 15 min.The 100μl of the resulting aqueous phase was transferred to a 96 well plate and 60 μl isopropanol added. The plate was placed on an orbital shaker at medium speed for 1 min. 20μl MagMaxTM beads were added to each well and plate returned to orbital shaker for 3 min. The 96 well plate was placed onto a magnetic separation rack and supernatant removed without disturbing the magnetic beads. This was repeated until all the aqueous phase had been used. The remaining extraction was performed as per the MagMaxTM −96 total RNA isolation manufacturers protocol, including a TURBOTM DNase clean up step, with a final total RNA elution volume of 40μl.

### Hippocampus whole tissue RNA extraction

100 mg of RNAlater-stabilised tissue was homogenised in 1 ml QIAzol reagent using a Qiagen TissueRuptor II on a medium speed setting for 40seconds, or until the lysate was uniformly homogeneous. Total RNA extraction was performed as per the Qiagen RNeasy Lipid tissue mini kit product guidelines, to a final elution volume of 40 μl.

### RNA quantification and quality control

Quantification and quality control of RNA samples was performed on an Agilent 4200 Tapestation. Tissue samples returned Total RNA with concentrations of 80-150ng/μl. Isolated cell suspensions returned total RNA with concentrations of 110-340pg/μl. A RIN cut-off was set at 6.0.

### RNA sequencing and analysis

Samples were prepared for sequencing using the Takara SMARTer stranded total RNA-Seq vs2 library prep protocol. Sequencing was performed as paired-end reads with a read length of 50bp. A total of 31 samples were analysed from 16 pigs (multiple brain regions were sampled for 12 pigs) comprising alveolar macrophages (n=3), microglia from cerebellum, frontal cortex, and hippocampus (n=8 per region), and hippocampal whole tissue (n=4). Raw sequence files (FASTQ format) were filtered and trimmed with BBtools, followed by alignment to the Sus scrofa genome (Sscrofa11.1; Ensembl Release 99) using Hisat2(Kim, Langmead, & Salzberg, 2015). Gene-level counts were generated from the resulting BAM files using StringTie (Pertea et al., 2015), and normalised gene expression (FPKM) data was subsequently made with Ballgown. Differential gene expression analysis was performed using Limma (Ritchie et al., 2015) on genes with > 10 CPM (counts per million) in all samples of at least 1 subgroup. TMM normalisation was used in EdgeR. Differential gene expression was carried out using Limma voom, using an adjusted p-value cut-off of 0.01. The fold change (FC) threshold was set to FC > 3 for defining microglial- and macrophage-enriched genes, and no threshold for characterising regional variations. The RNA-seq data is available via Gene Expression Omnibus (GEO https://www.ncbi.nlm.nih.gov/geo/, accession number: GSE172284).

Gene expression in microglia isolated from frontal cortex, cerebella and hippocampus was compared to those in macrophages, and from this, macrophage-enriched genes were defined. The microglia enriched genes were derived using genes found to be more highly expressed in microglia relative to both macrophage and corresponding tissue (hippocampal microglia compared to hippocampus tissue). A microglial gene list was generated by refining the common DEG from the aforementioned gene expression comparisons through the use of mouse and human brain expression data from www.brainrnaseq.org (Bennett et al., 2016); genes were removed if they were expressed in nonmicroglia cell types (> 2 FPKM) and were less than 8-fold higher in microglia than the non-microglia cell types in both the human and mice datasets. As some pig genes were not annotated with a gene name, human orthologues were used in addition to the pig gene names.

When examining regional variations, each pair-wise comparison (cerebellum vs frontal cortex, frontal cortex vs hippocampus, hippocampus vs cerebellum) was carried out. For each of the genes showing significant regional variation, the different brain regions were ranked according to their gene expression level and the region with the highest expression noted.

Gene-gene co-expression network analysis was carried out using data for microglia isolated from cerebellum, frontal cortex and hippocampus using Graphia Professional (version 2.1. Kajeka, Edinburgh, UK), with a Pearson correlation threshold of >0.84 on genes with > 1FPKM in at least 1 sample, followed by Markov clustering (MCL) using an inflation value of 1.8. Gene Ontology (GO) enrichment analysis was carried out on both genes showing significant regional variations and annotations in the Pig Expression Atlas (Freeman et al., 2012); an adjusted p-value < 1 × 10^−5^ and a minimum of 10 genes were required for an enrichment to be considered significant. We carried out the co-expression analysis for two reasons. Firstly, it is more inclusive by including additional genes beyond those meeting statistical thresholds based only on pair-wise filtering, thereby allowing larger gene sets to be used in the GO enrichment analysis. Secondly, if multiple processes were associated with a brain region, the genes involved in each process can be deconvolved, allowing for better defined input for GO enrichment analysis.

Sample-sample network analysis examining the relationships of microglia isolated from the three brain regions was carried out in Graphia with log2 FPKM and a Pearson correlation coefficient threshold > 0.85 on genes showing significant regional variation, with MCLi set to 1.8.

Functional enrichment was carried out using Metascape (Zhou et al., 2019), with the settings for min overlap, p value cutoff and min enrichment set to 3, 0.01 and 1.5 respectively, using the Gene Ontology (GO) Biological Processes database.

### Cross-species comparison of microglial transcriptome signatures

Raw RNAseq data on isolated microglia from mice (n=17) were obtained from Geirsdottir et al. (2019) (NCBI BioProject: PRJNA556201). Non-pathogenic human microglia RNBAseq data were from 2 separate studies (n=6; NCBI run IDs: SRR9909238, SRR9909237, SRR9909236, SRR6849268, SRR6849266 and SRR6849267) (Sankowski et al., 2019; van der Poel et al., 2019). These data were integrated with six of the pig microglia samples from this study (two from each brain region). Processing of data from raw FASTQ files to FPKM were the same as those used in processing the pig data for this study. Reference genomes and gene annotations for the respective species were downloaded from the Ensembl databases (Release 99). For each species, the orthologue genes matching to each human gene were noted using Ensembl Biomart (Ensembl Genes 99) (Kinsella et al., 2011). In cases where multiple orthologue genes mapped to the same human gene, the sum FPKM were used. We created a merged and normalised RNAseq dataset that contained annotated genes mapping across the three species using homologues that matched to the corresponding human gene in the Biomart database (GRCh38.p13).

Analysis was limited to genes with matching human homologues across all species. These genes were ranked according to their FPKM, followed by minmax normalisation of their ranks to a value between 0 and 1, with 0 indicating the least expression and 1 the highest expression. Following this, the expression of microglia-enriched genes was examined for four gene lists: pig microglia-enriched genes (derived from this study), human microglia-enriched genes [two gene lists from (Galatro et al., 2017; Patir et al., 2019) and mouse microglia-enriched genes (Butovsky et al., 2014). Two human microglia-enriched gene lists were used to cover different methods deriving an enriched microglial gene signature; one from co-expression (Patir et al., 2019) and the other from fold-change enrichment (Galatro et al., 2017). For the gene list from Galatro et al. (2017), we included the additional filter on the microglia-gene list provided in the manuscript, keeping only genes with adjusted p < 0.001 and FC>2 when comparing microglia to monocytes, and macrophages.

## Results

### Identification of comparative pig microglia and macrophage transcriptomes

We first defined the pig microglia-enriched gene set by comparing the transcriptome of sorted microglia with isolated alveolar macrophages (as an exemplar systemic macrophage comparator) and whole hippocampal tissue from the same animals. The pattern of expression for selected canonical genes enriched in microglia, other parenchymal CNS cell types, and CNS border/systemic macrophages provided initial validation of the specificity of the cell sorting procedure (CD11b^+^CD45^lo^ for microglia, F4/80^+^ for alveolar macrophages, Figure S1). Notably, microglial samples expressed high levels of genes established as enriched in microglia in other species and negligible levels of CNS border macrophage-enriched genes (e.g. CD163, MRC1) that, in contrast, showed a reciprocal pattern in alveolar macrophages (Figure 1). Expression of archetypal neuronal, astrocytic, oligodendroglial and vascular genes were also negligible in sorted microglial samples (Figure 1). These data confirm specificity of the cell sorting protocol for parenchymal microglia.

**Figure 1.**
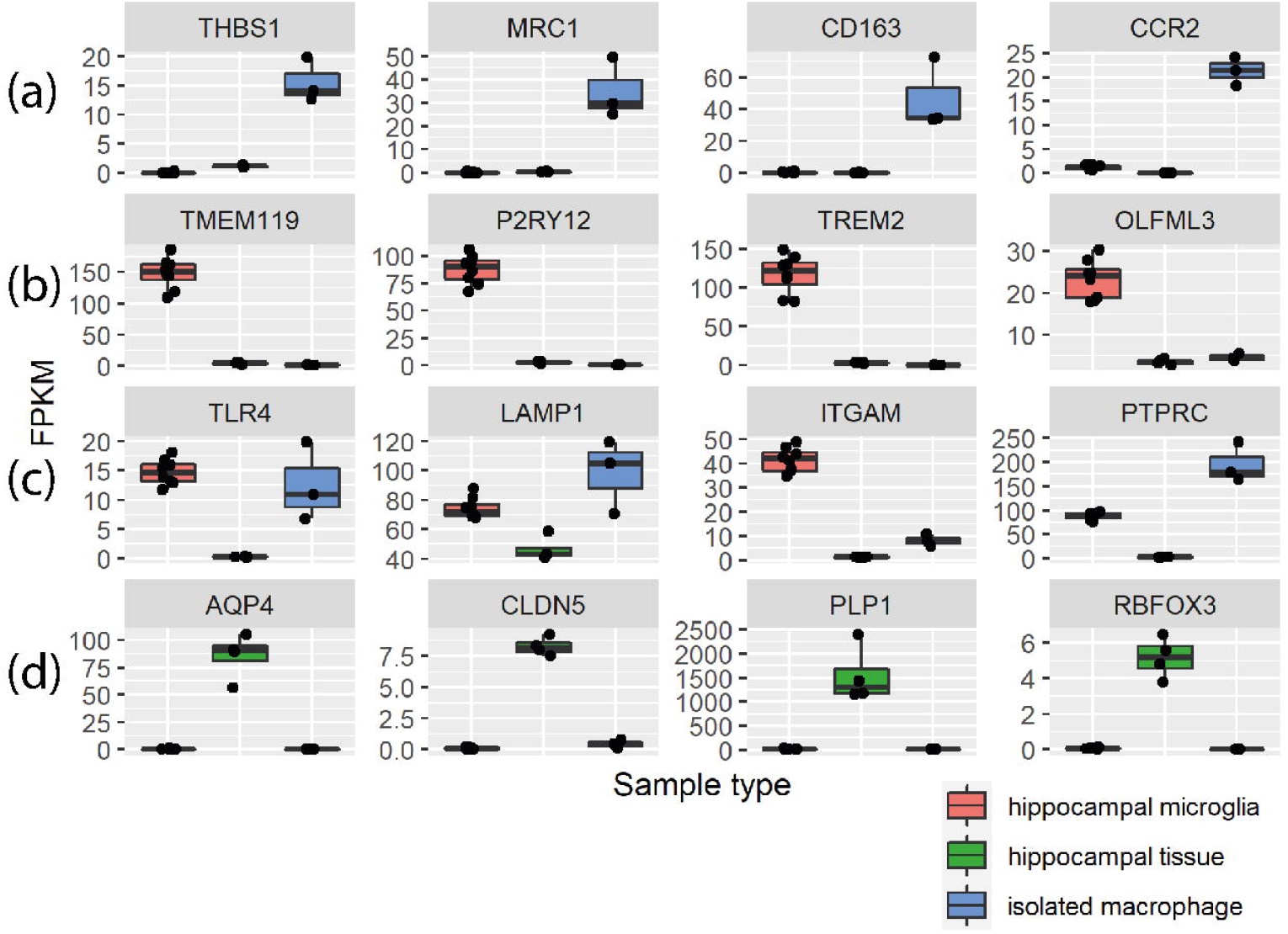
Specificity of cell isolation according to selective expression of canonical cell-type genes in isolated cells and brain tissue. Expression of genes in our RNAseq dataset for established enriched genes from previous studies in rodents or humans for (a) non-CNS monocytes/macrophages, (b) microglia, (c) myelomonocytic cells, (d) CNS cell types (AQP4, astrocytes; CLDN5, endothelial cells; PLP1, oligodendroglia, RBFOX3, neurons).

Differentially expressed genes (DEG, (adjusted p < 0.01, FC > 3) were used to define pig microglial signature genes, which are genes with high expression in microglia compared to alveolar macrophages and whole brain extracts from the hippocampus. A total of 959 genes were more highly expressed (Table S1) in microglia versus macrophages, and 1,586 genes were more highly expressed in isolated hippocampal microglia compared to hippocampal whole tissue (Table S2). Amongst these sets of DEGs, 499 were present in both comparisons. Although we confirmed high specificity of microglial sorts (see above) we reasoned that even a very minor contamination with other CNS cellular constituents could result in enrichment when compared to alveolar macrophages and potential aberrant inclusion in the microglial signature. We therefore refined the 499 gene list by cross-checking (see Methods) expression against all cell types in the Brain RNAseq dataset (http://www.brainrnaseq.org) (Bennett et al., 2016) (Table S3).This created a highly stringent set of 365 genes comprising the pig microglia-enriched gene list (Table S4 and S5, Figure 2). We observed that many genes previously shown to be microglial-enriched genes across other species were also contained in the porcine list (e.g. *C3, CSF1R, CX3CR1, GPR34, OLFML3, P2RY12, TREM2*).

**Figure 2.**
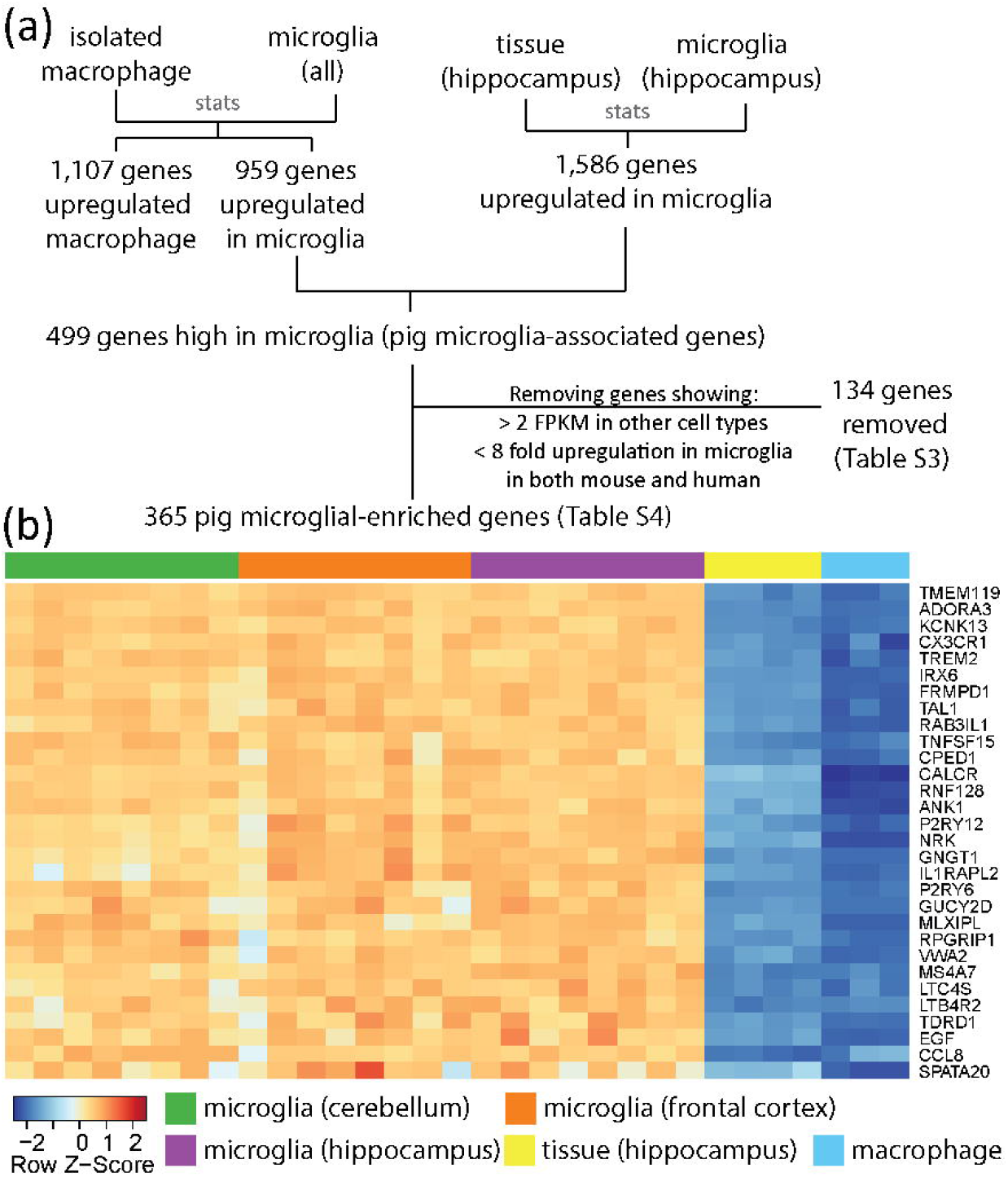
Derivation of the pig microglia gene expression profile. (a) The 959 upregulated differentially expressed genes (DEG) when comparing isolated microglia to macrophages have 499 genes in common with the 1,586 upregulated DEGs when comparing isolated hippocampal microglia to hippocampal tissue. By using data from Brain RNAseq (https://www.brainrnaseq.org/), we excluded 134 genes (Table S3) showing < 8 fold upregulation in microglia and >2 FPKM expression (Fragments Per Kilobase of transcript per Million) in human or mouse astrocytes, neurons, oligodendrocytes or endothelial cells from the 499 DEGs, yielding a final 365 genes as the pig microglia-enriched genes (Table S4). (b) The top 30 most enriched genes in microglia (with gene names) are shown in the heatmap (Table S5). Z-score was used to represent the FPKM expression as standard deviations away from the mean for each gene.

We also assessed the genes that were enriched in macrophages compared to microglia to further understand the differential gene signatures. 1,107 genes were significantly more highly expressed (adjusted p < 0.01, FC> 3) in isolated alveolar macrophages compared to microglia. We further filtered these (see Methods) to minimise inclusion of genes that might arise from even negligible numbers of contaminating immune cells (e.g. T cells, B cells) in the alveolar samples by surveying expression across the major immune cell classes in the ImSig dataset (Nirmal et al., 2018) (Table S6). This produced a high-stringency set of 1,032 genes enriched in macrophages compared to microglia (Table S7, Figure 3). The top 30 macrophage-enriched genes (highest FC and with gene annotation) were are all expressed at < 1 FPKM in microglia samples (Figure 3, Table S8) showing they are both enriched for macrophages relative to microglia and indicating they could be a useful set of negative selection markers for studying pig microglia. In summary, we have created high-stringency pig microglial and systemic (alveolar) macrophage gene signatures (Table S4) highlighting the distinct identities of these tissue macrophage populations (Table S7).

**Figure 3.**
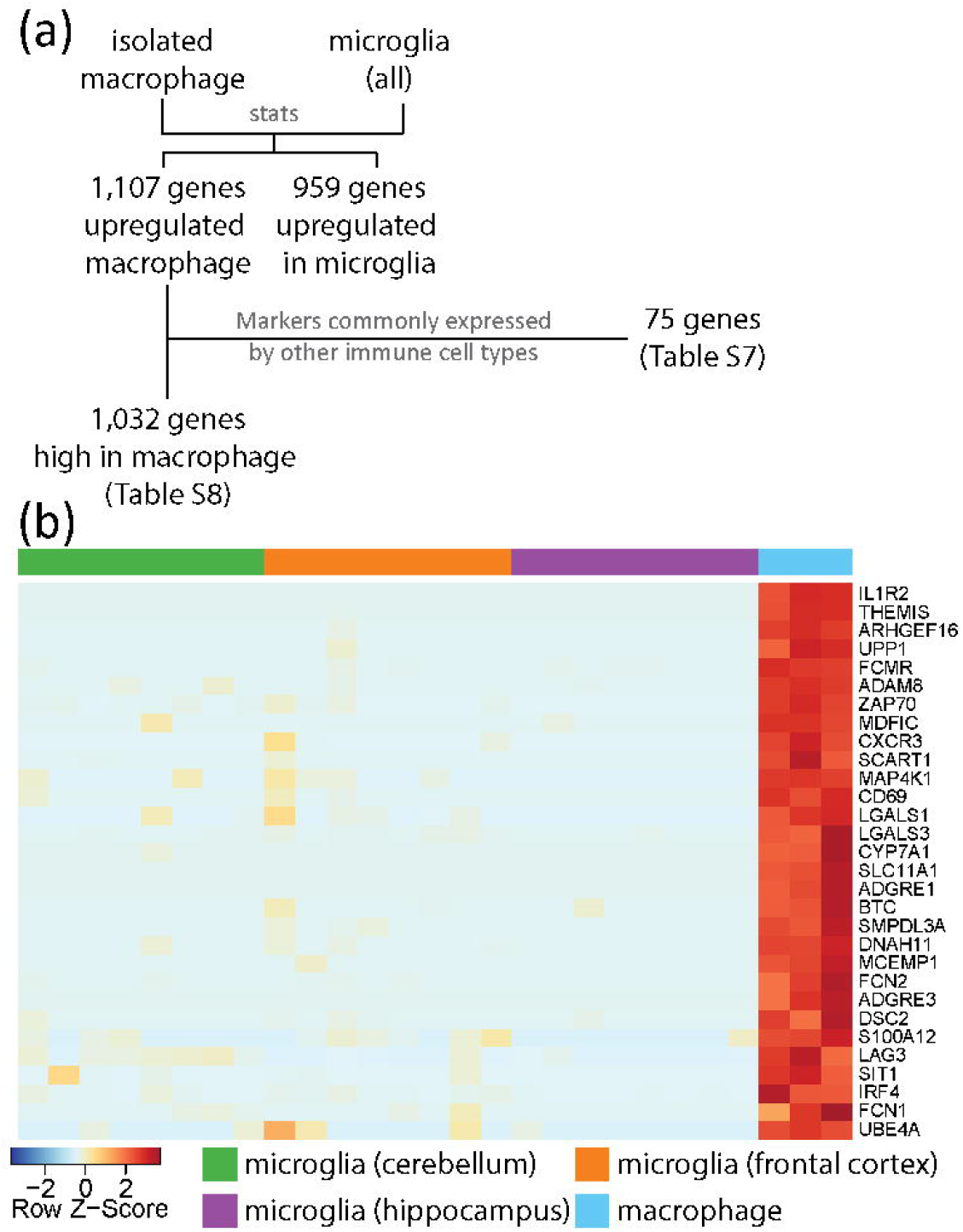
Genes enriched in isolated macrophages compared to microglia. (a) When comparing isolated macrophages to microglia, 1,107 genes are upregulated in macrophages. Of these, 75 genes were markers commonly expressed by other immune cell types such as T-cell, B-cell, and plasma cells [48 genes in ImSig (Nirmal et al., 2018), 27 genes are related to immunoglobulin; e.g. CD2, PVRIG, CD19, NLRC3, CD8A], these are highlighted in Table S6. The remaining 1,032 gene can be found in Table S7. (b) The heatmap illustrate the top 30 genes (with gene names) showing the highest enrichment in macrophages (Table S8). Z-score was used to represent the FPKM expression as standard deviations away from the mean for each gene.

### Regional variation of the pig microglial transcriptome

Studies in mice and humans have reported differences in the transcriptional profiles of microglia from distinct brain regions (Grabert et al., 2016; Hammond et al., 2019; Sankowski et al., 2019). We therefore determined whether the microglia in the pig brain showed regional diversity by comparing microglia from the cerebellum, hippocampus and frontal cortex. This analysis identified 215 genes with significant variation across these regions (adjusted p < 0.01; Table S9). Amongst the 215 regionally variant genes, only 24 genes overlapped with the 365 pig microglia signature genes (Figure 4c and Table S10; e.g. *LPAR5, CD1D, MS4A7, C3, FBP1*, and *SLC2A5*), suggesting that differential expression of core identity genes contributes relatively little to regional heterogeneity. Using these genes to carry out sample-sample correlation analysis suggested that cerebellar microglia were particularly distinct from the microglia in the hippocampus and frontal cortex (Figure 4a). Of the regionally-variant genes, 121 were most highly expressed by cerebellar microglia, 70 genes were most highly expressed by frontal cortex microglia, and 24 were genes most highly expressed by hippocampal microglia (Figure 4b). The intermediate profile of hippocampal microglia evident from sample-to-sample correlation and differential gene expression was similar to previous findings in rodent brain and indicative of a comparable rostro-caudal gradient in profile (Grabert et al., 2016). GO enrichment on the 215 genes showing regional variation indicated enrichment in genes involved in extracellular structure organisation (e.g. GO:0043062, p < 10^−8^, Figure S2).

**Figure 4.**
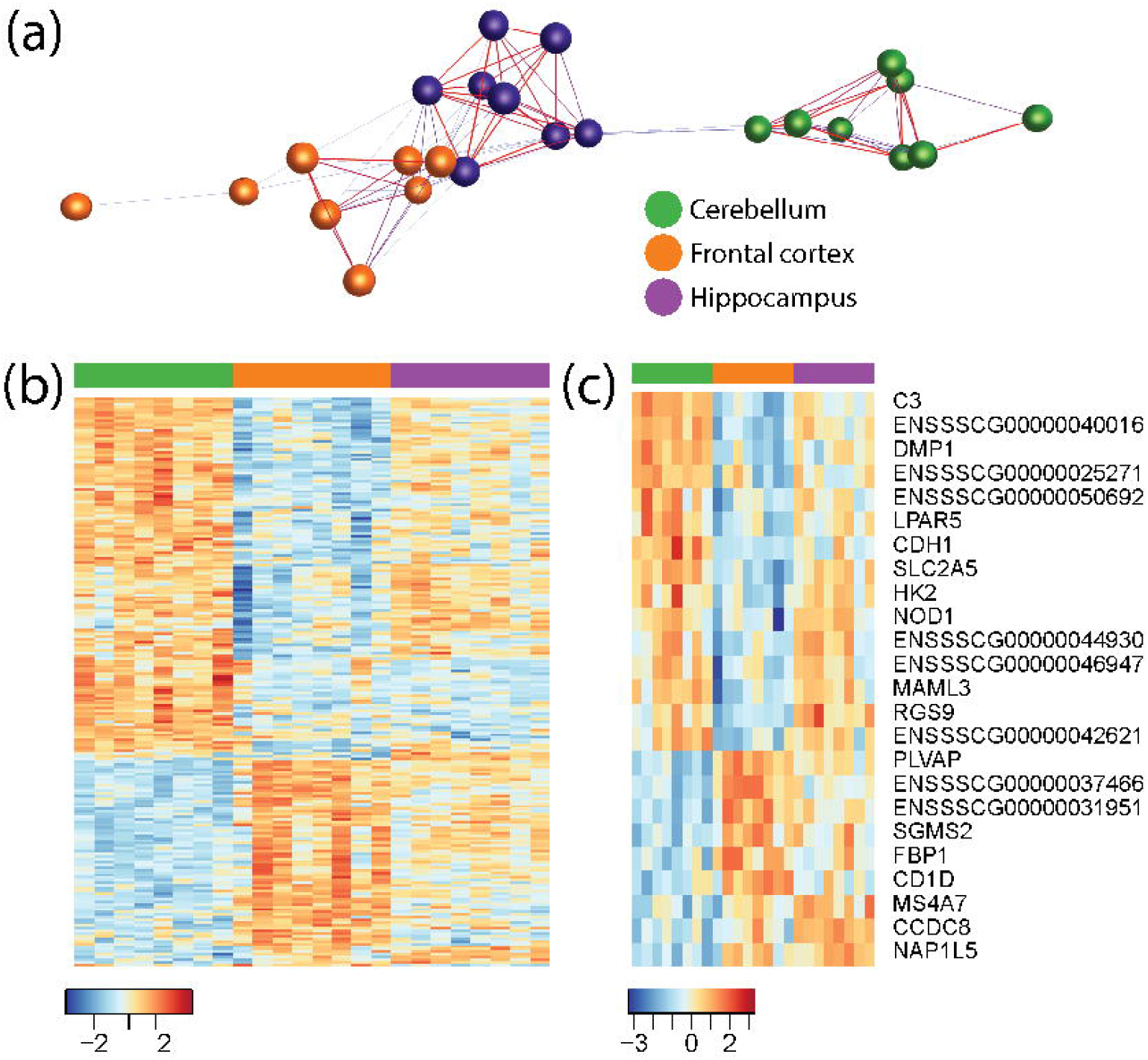
Regional variation of the pig microglial transcriptome. The gene expression for microglia from the frontal cortex, hippocampus and cerebellum were compared to each other, and 215 genes show significant regional variation (Table S9). (a) Using the 215 genes showing high regional variation for sample-sample correlation analysis, microglia from the frontal cortex (orange) and the hippocampus (purple) are clustered closer to each other than to those from the cerebellum (green). (b) Heatmap illustrating the differential expression of the 215 genes across microglia from the three regions. (c) When comparing these 215 genes showing regional variation against the 365 pig microglia-associated genes, 24 genes are present in both gene lists. Z-score was used to represent the FPKM expression as standard deviations away from the mean for each gene.

To explore in more depth the biological processes associated with specific regional microglial variation, we examined gene clusters enriched for the 215 genes showing regional-variation from a co-expression gene network comprising microglia from the cerebellum, hippocampus and frontal (r=0.84, MCLi=1.8) (Table S11). The co-expression network analysis highlighted three co-expressed gene clusters that were enriched in the 215 regionally variant genes (Figure 5, Table S11): Cluster 5 (lower in cerebellum; e.g. *PDCD4, ACSL3, SNX3*, and *ARPC5*); Cluster 19 (high in cerebellum; e.g. *CDH1, CSF1*, and *SLC15A3*); Cluster 37 (high in cerebellum; e.g. *C3, BCL6*, and *BTG1*) (Table S11). Cluster 5 genes (relatively higher in frontal cortex and hippocampus) were significantly enriched in GO processes associated with myeloid cell activity and metabolism, including myeloid leukocyte activation (e.g. GO:0002274, p<10^−7^, Figure S3, e.g. *AIF1, ARPC5, CD63, FCER1G, RHOA*), oxidative phosphorylation, phagocytosis, and antigen processing and presentation. Genes in cluster 19 and 37 (relatively higher in cerebellum) were also significantly enriched in GO processes associated with myeloid leukocyte activation (e.g. GO:0050920, p < 10^−6^, Figure S4, e.g. *C5AR2, CSF1, CXCL14, EGR2*), and also in regulation of the adaptive immune response (GO:0002822, p<10^−4^, Figure S5, e.g. *SMAD7, BCL6, C3*) and cell adhesion/migration/chemotaxis).

**Figure 5.**
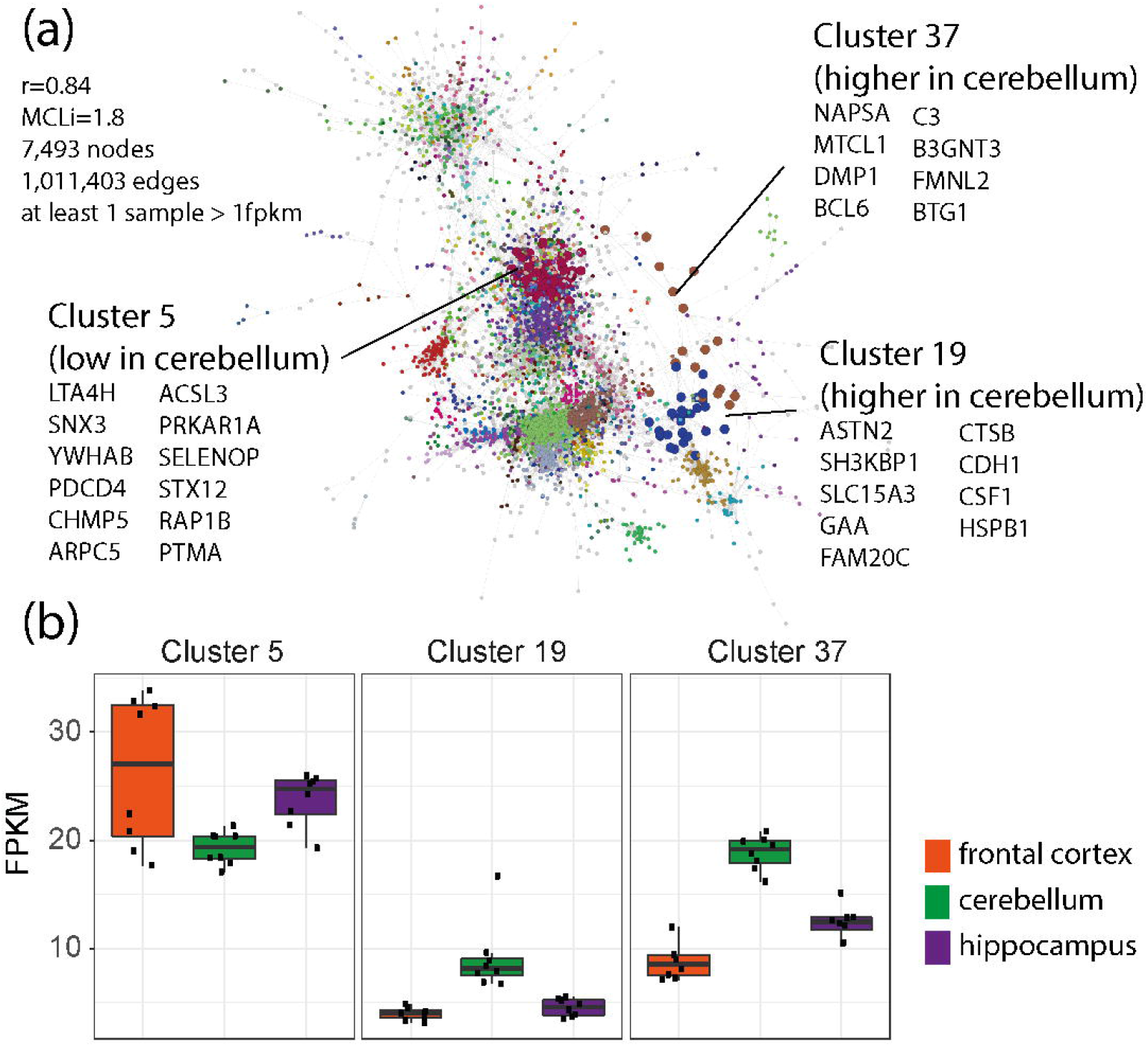
Gene-gene co-expression network analysis of microglial samples. Co-expression network analysis was carried out using all pig microglia samples (Table S11), (a) The co-expression clustering method groups together genes showing similar pattern of expression across samples. Three clusters are found to be enriched in genes showing significant regional variation, Cluster 5, 19 and 37. (b) The average expression levels (FPKM) for each of the clusters are shown in the figure (y-axis), and each bar represent an individual microglia sample from frontal cortex (orange), cerebellum (green) or hippocampus (purple). Cluster 19 (such as CSF1, CTSB, CDH1) and 37 (such as C3, DMP1, NAPSA) are higher in expression in the cerebellum, whilst cluster 5 (such as ACSL3, PDCD4, RAP1B) is lower in cerebellum.

### Cross-species comparison of microglial transcriptional signatures

We conducted a comparative analysis of the pig microglial transcriptome profile identified here with those in mice and humans. A total of 15,199 genes with a matching human homologue across mice, humans and pigs were identified. To allow cross-dataset and cross-species comparison, the RNAseq data were expressed as FPKM and normalised to percentile expression, with 1 being the maximum expression and 0 being the minimum expression (Figure 6). Sample-sample correlation was then carried out on the normalised combined dataset containing the 15,199 genes. This analysis revealed that the pig and human microglia samples clustered closely to each other and separate from the mouse microglia in the network graph (Figure 6).

**Figure 6.**
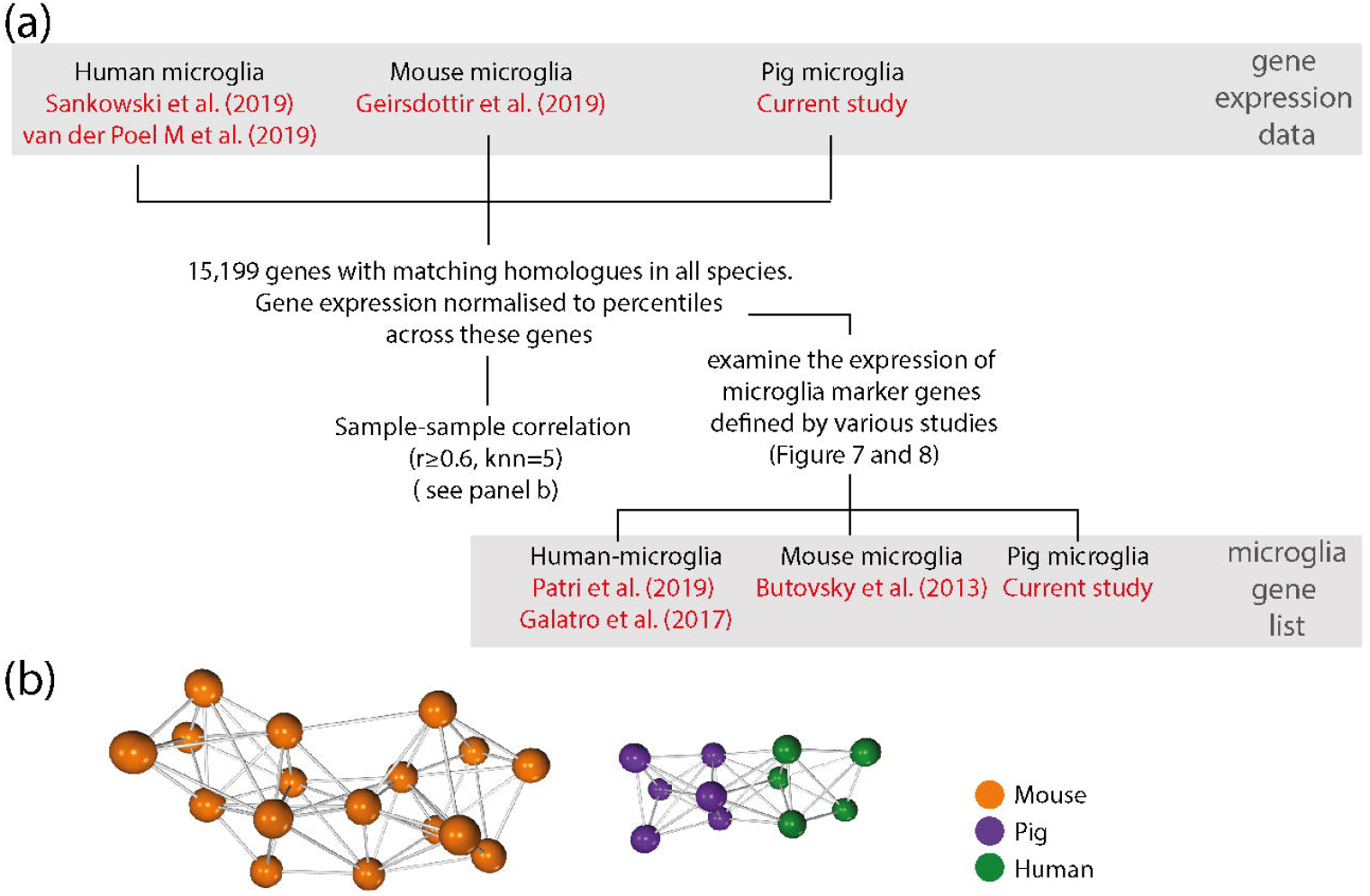
Workflow to compare human, pig and mouse microglia transcriptomes. (a) Genes with homologues across human, pig and mouse were selected, a total of 15,199 genes. In order to create a combined dataset that can be used to compare across species and data sources, the expression of genes for each sample was normalised to the percentile of expression for these 15,199 genes. Subsets of genes were used to further examine the species-variation in human microglia-associated gene expression. The expression of human, mouse and pig microglia marker genes were examined across these datasets. Two human microglia gene lists were included, one derived from Patir et al. (2019) and the other from Galatro et al. (2017); the former was based on coexpression, while the latter was based on differential expression. The mouse microglia gene list was derived from Butovsky et al. (2014), and the pig microglia gene list from the current study. With the exception of the pig microglia, the gene expression data and the gene lists were from separate studies. (b) Sample-sample correlation using the normalised dataset suggests that overall, human microglia are more similar to pig microglia than those of mouse.

We further explored microglial species relatedness by comparing the expression of microglia signatures gene profiles derived from pigs (this study), human (Galatro et al., 2017; Patir et al., 2019) or mouse (Butovsky et al., 2014) across the cross-species dataset of 15,199 genes described above (Figure 7 and 8). K-means clustering was used to group these genes into 5 clusters for each gene list (Figure 7 and 8 and Table S12-15). As was expected, for each microglial species-derived signature gene list, the expression of those signature genes is highest in samples of the same species on the cross-species dataset (cluster 5 in Figure 7; example genes shown in Figure 8). This is evident even when the signature gene list and expression data on which the gene list is mapped are derived from different studies. One exception is a small cluster of mouse microglia signature genes (derived from Butovsky et al. 2014) that is not highly expressed by the mouse microglia from Geirsdottir et al. (Cluster 3 in Figure 7c); these may indicate study-specific genes.

**Figure 7.**
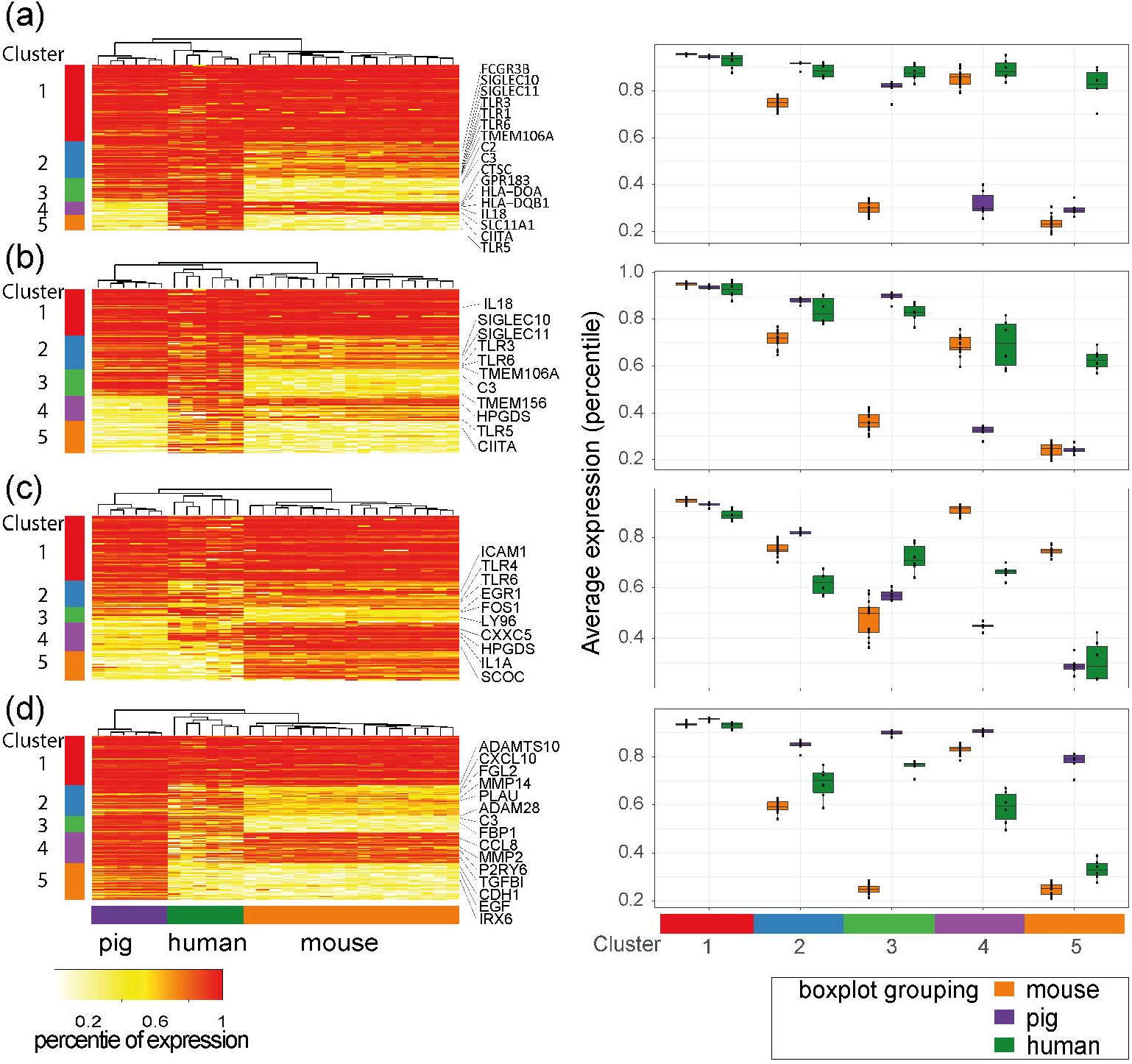
Inter-species comparison of microglial core signatures. Gene expression data were compared across studies by converting FPKM to percentile-expression. We then examined the expression of human, mouse and pig microglia gene lists in the gene expression data for the three species. By using k-means clustering, 5 groups of genes are characterised for each gene list (cluster 1-5), and the average expression for each cluster is shown per species on the right hand side of the corresponding heatmaps. (a) Of the human-microglia genes reported by Patir et al. (2019), 170 genes have matching orthologues in the combined dataset. Cluster 1 is expressed in all three species, whereas Cluster 2 and 3 are less expressed in mouse microglia. Cluster 4 is low in pig and cluster 5 being low in both pig and mouse. (b) Of the 252 human microglia genes reported in Galatrox et al. (2017) (additional filters described in methods, 156 genes have matching orthologues in the combined dataset. The observed pattern is similar to analysis done with human microglia markers defined through co-expression in panel a. (c) Of the 152 mouse microglia genes reported by Butovskyet et al. (2014), 126 genes have matching orthologues in the combined dataset. Cluster 1 is expressed in all three species, whereas Cluster 2 is slightly less expressed in human. Cluster 3 is less expressed in mouse microglia. Cluster 4 is low in pig, and cluster 5 is low in both human and pig. (d) Of the 365 pig microglia genes reported in this study, 179 genes have matching orthologues in the combined dataset. Cluster 1 is expressed in all three species, whereas Cluster 2 and 3 are slightly less expressed in mouse. Cluster 4 is expressed in all species, though to a lesser extend in human. Cluster 5 is low in both human and mouse.

**Figure 8.**
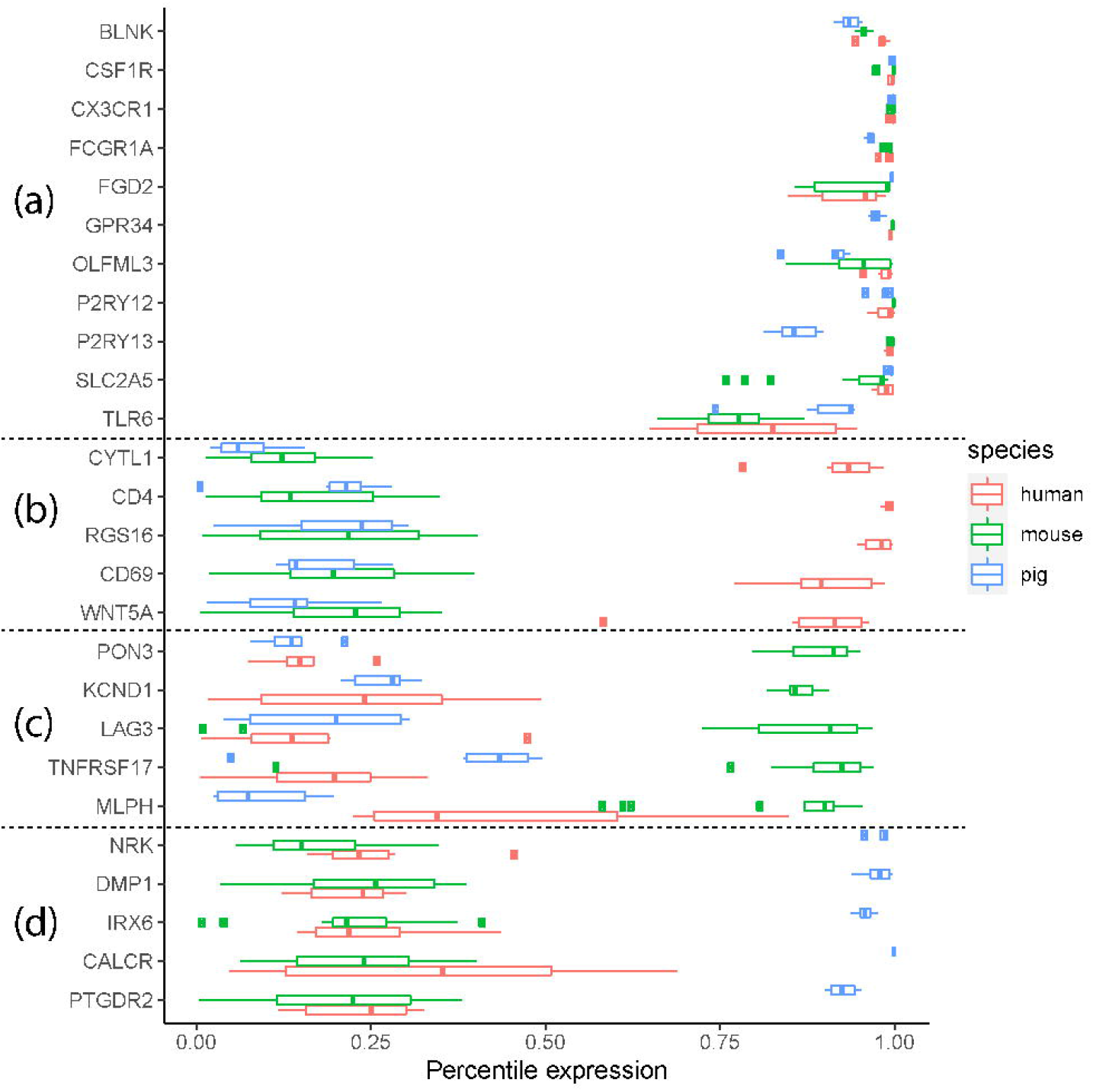
Examples of species-common and species dependent microglia gene expression. A selection of the species common or species unique genes and their expression are shown for (a) genes present in Cluster 1 for all gene lists, (b) gene present in human-microglia-unique cluster in Figure 7a or 7b, (c) genes present in mouse-unique cluster in Figure7c, and (d) genes present in the pig-microglia-unique cluster in Figure 7d.

For all species gene signature lists, there is a cluster of genes that is highly and similarly expressed across all species (labelled as Cluster 1 in each list). There is a total of 138 unique genes after combining all genes from Cluster 1 of each study (Table S12-15), 10 of which are present in all 4 gene lists (*BLNK, CSF1R, CX3CR1, FCGR1A, FGD2, GPR34, OLFML3, P2RY12, P2RY13, SLC2A5*), which we note include several archetypal microglial signature genes, reaffirming their conserved nature. The remaining clusters demonstrated species-related patterns of expression. Clusters 2 and 3 commonly showed similarity between pig and human microglia whereas cluster 4 showed greater similarity between mouse and human microglia. Cluster 5 was highly species-selective with genes expressed highly in the cross-species expression dataset from samples corresponding to the species from which the gene signature was derived, as expected. Overall, we detected a greater number of clusters and genes showing a more similar level of expression between pig and human microglia than between mouse and human microglia and is reflected in the hierarchical clustering of samples when using the human- and mouse-derived microglial signature gene lists. This is consistent with and adds to the sample-to-sample correlation network analysis above (Figure 6) showing closer relatedness for pig with human microglia.

## Discussion

Here, we conducted the first study that defines the pig microglial transcriptome-wide gene signature and show regional variation in pig microglial gene expression. Moreover, we showed that for a portion of the human microglial markers, pig microglia have a gene expression pattern more similar to human microglia than mouse microglia, reinforcing the utility of the pig as a translational and complementary species for the study of microglia and neuroinflammation in disease and other challenges to brain homeostasis such as psychosocial stress.

We have reported 365 genes highly enriched in isolated pig microglia relative to macrophages and whole brain tissue from the same brain region. Many of the top 30 pig microglia genes identified in this study have previously been noted as amongst the most enriched genes when comparing microglia to monocytes/macrophages in multiple mouse genome-wide gene expression studies, including *P2RY12, TMEM119, TREM2*, and *CX3CR1* (Butovsky et al., 2014; Chiu et al., 2013; S. E. Hickman et al., 2013). *LTC4S* has been validated to be microglia-enriched in mice (Bennett et al., 2016). SALL1 is not only uniquely expressed by microglia (among adult CNS cell types), but has been found to play a role in the maintenance of microglial identity as its inactivation converts microglia into inflammatory phagocytes (Buttgereit et al., 2016). *TREM2, TMEM119, CX3CR1* and *MLXIPL* have also been reported from human studies to be increased in microglia compared to monocytes (Butovsky et al., 2014). *P2RY6* has been demonstrated to be involved in microglial phagocytosis in rat microglia *in vitro* (Koizumi et al., 2007). *RAB3IL1* and *LTC4S* are found in the full gene list from various human or mice studies examining microglia gene signatures (Bennett et al., 2016; Butovsky et al., 2014; Patir et al., 2019). A recent study showed conservation of a core microglial signature across multiple species (Geirsdottir et al., 2019), and although this did not analyse the pig microglial transcriptome, our data are consistent with the concept that microglia, including in the pig, have core features that govern conserved key functions across evolutionary diverse mammalian species.

We also examined the variation in gene expression by microglia from different brain regions of the pigs. In additional to acting as immune sentinels, microglia also play also important roles in CNS homeostasis during development and in adult health and disease (Prinz & Priller, 2014). To carry out these multifunctional roles, microglia are required to sense perturbations in their environment to elicit appropriate microglial responses to maintain homeostasis (Grabert et al., 2016; Stratoulias, Venero, Tremblay, & Joseph, 2019). For instance, exposing microglia to signals by healthy neurons appears to promote their resting state and antagonize pro-inflammatory activities (Biber, Neumann, Inoue, & Boddeke, 2007). Fractalkine (CX3CR1-CX3CL1) signalling has been found to regulate microglial activation; neuronal membrane bound CX3CL1 maintains CX3CR1-expressing microglia in a surveying state, and cleaved soluble CX3CL1 is thought to stimulate migration of inflammatory cells (Szepesi, Manouchehrian, Bachiller, & Deierborg, 2018). The mammalian brain is organised into regions with specific biological functions and properties with distinct transcriptional and metabolic profiles (Choi et al., 2018; Hawrylycz et al., 2012). Indeed, microglial regional variation has been described not only in their distribution and morphology (Lawson, Perry, Dri, & Gordon, 1990; Savchenko, Nikonenko, Skibo, & McKanna, 1997; Tan, Yuan, & Tian, 2020), but also in their gene and protein expression in both mice and human (Bottcher et al., 2019; Grabert et al., 2016; Patir et al., 2019; Sjostedt et al., 2020), although the extent of this may depend on the analytical method used. Our study has observed 215 microglial genes showing regional heterogeneity in the pig brain, with 24 such genes also being microglial markers. In this study, we found microglia from the cerebellum to be more distinct from their counterparts from the frontal cortex and hippocampus, which is an observation reported by other studies (Grabert et al., 2016; Soreq et al., 2017). There is a higher level of expression in genes related to cell-substrate adhesion in microglia derived from the cerebellum. Cerebellar, but not striatal or cortical, microglia have been shown to display unique high levels of cell clearance activity (Ayata et al., 2018). The finding that only 10% of regionally heterogenous genes are also part of the core signature gene set is similar to our previous observations in the mouse brain and indicates that regional heterogeneity is primarily superimposed upon core identity (Grabert et al., 2016). Nonetheless, since 24 of these 215 regionally varied genes were also present within our core pig microglia signature gene list, it is important to consider that the expression level of some core genes may vary across the brain when compared to the others and selection of appropriate markers (e.g. for labelling microglia) may require consideration of the regions being analysed.

When comparing the expression of microglial gene signatures derived from human, mouse and the present pig datasets, there is a group of microglial genes that are highly expressed by microglia of all three species. Whilst in phylogenetic terms rodents are more closely related to humans than pigs to humans (Song, Liu, Edwards, & Wu, 2012), we have found that pig microglia have a more similar expression pattern to humans than mice on a transcriptome-wide scale and for expression of microglial core signature genes. There may be elements of convergent evolution perhaps relating to brain and organismal size, adaptations to environment, and social behaviours, that contribute to inter-species relatedness in anatomical and physiological features. Brain mass/volume, grey-white matter composition and neuronal densities and size are all more similar between humans and pigs and may affect glial characteristics (Herculano-Houzel, 2014). Geirsdottir et al. (2019) have noted complement genes are expressed at a lower level in rodent microglia than human microglia. Our study has demonstrated that pig microglia have higher levels of expression of complement components *C2* and *C3*, more similar to human microglia than mouse microglia. Genes in the complement system, such as *C3*, have previously been demonstrated to play a central role in Alzheimer’s disease (Lian et al., 2016). Galatro et al. (2017) have highlighted several immune genes such as *TLR, Fc*, and *SIGLEC* receptors, to be abundantly expressed in human microglia but not in mouse microglia. When comparing pig, human and mouse microglia, we found groups of genes showing higher level of expression in human and pig microglia compared to the mouse, including *TLR1, TLR3, TLR6, FCGR3B, SIGLEC10* and *SIGLEC11*. TLR3 expression has been positively correlated with plaques in Alzheimer’s disease as well as colocalising with the phagocytic marker CD68 (Walker, Tang, & Lue, 2018). Although more human microglia markers are expressed by pig microglia than mouse microglia, there is a subset of genes that is more similar between mouse and human microglia, and each species exhibit species-specific markers. Genes showing lower expression in pigs include *HLA-DQA, HLA-DQB1, SLC11A1* and *CTSC*. It is possible that unavoidable differences in methods for microglial isolation in different species/studies may influence the extent of species microglial relatedness, however our pig isolation protocol is similar to commonly used rodent protocols and we saw similar cross-species patterns when the human microglial signature was derived from different studies using distinct methods. Noting the differences as well as the similarities in gene expression pattern across microglia from different species highlights the importance of characterising pig-specific microglial signatures for facilitating a better understanding of pig neuroimmunology and pathology, and utility of the pig in translational biomedical and agricultural research.

In conclusion, we have defined the pig microglia transcriptome signature that distinguishes microglia from other CNS cell types and non-CNS macrophages, proposing gene sets that can be used for differentiating the different myeloid cell types in the pig. We have demonstrated regional variation in pig microglial gene expression, with those derived from the cerebellum being more distinct from those from the frontal cortex and hippocampus. Our results indicate that pig microglia and human microglia show a more similar gene expression pattern than mouse microglia and human microglia.

## Acknowledgment

This work was funded by a Wellcome Trust-University of Edinburgh Institutional Strategic Support Fund (Reference IS3-R77). Barry McColl acknowledges UK Dementia Research Institute support; UKDRI receives its funding from DRI Ltd, funded by the UK Medical Research Council, Alzheimer’s Society, and Alzheimer’s Research UK. Neil Mabbott, Barbara Shih, Sarah Brown and Alistair Lawrence acknowledge Roslin Institute strategic grant funding from the U.K. Biotechnology and Biological Sciences Research Council (grants BBS/E/D/10002071, BBS/E/D/20002173, BBS/E/D/20002174, BBS/E/D/3000227). Alistair Lawrence also receives funding support from the Scottish Government’s Rural and Environment Science and Analytical Services Division (RESAS).

