## Supplementary material for "Defining the pig microglial transcriptome reveals their core signature, regional heterogeneity, and similarity with humans": Figure S1

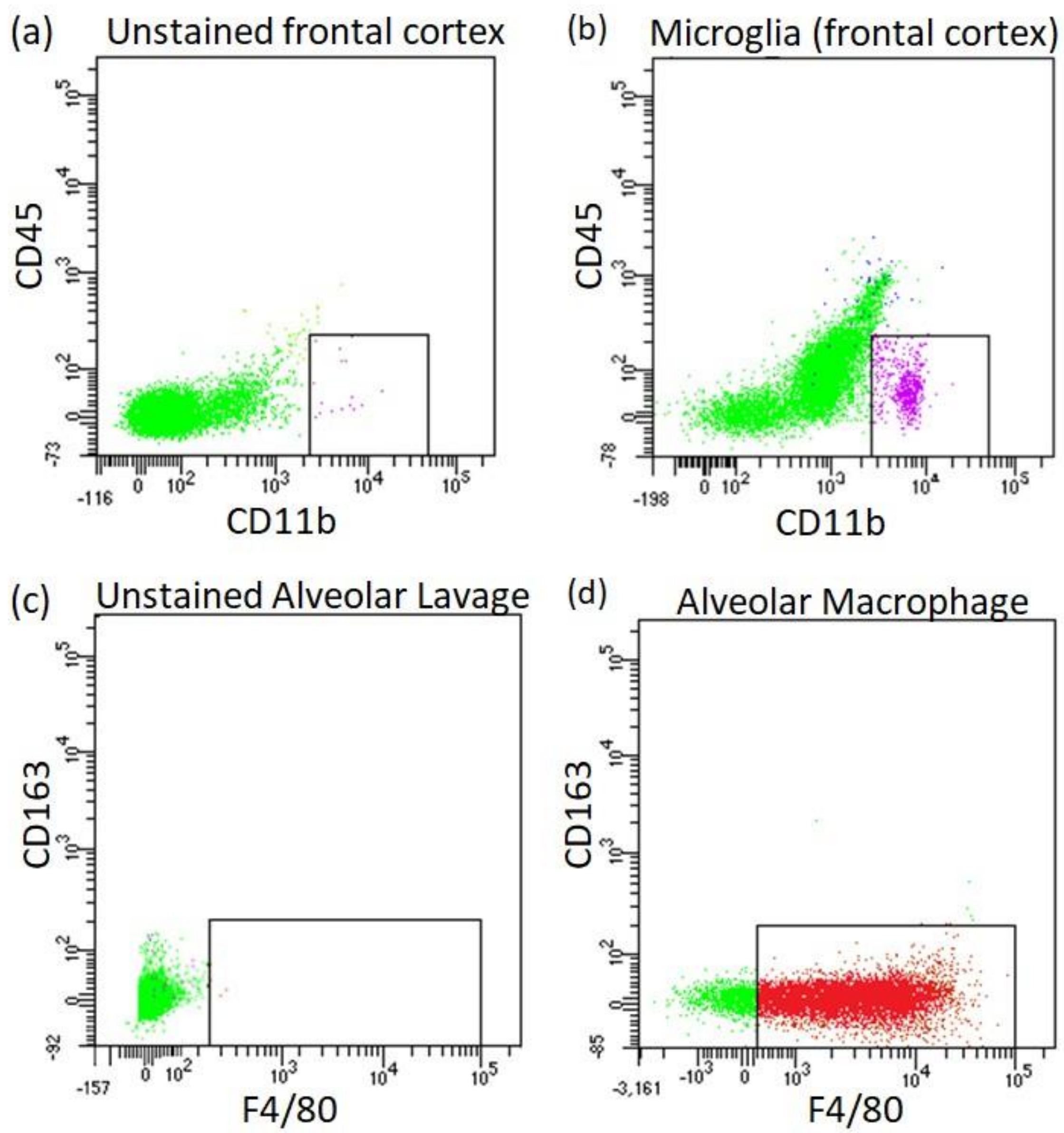


Figure S1. FACS gating for cell isolation

Flow cytometry of single cell suspensions derived from brain or lung. Each plot shows the fluorescence intensities of events corresponding to the fluorophore on the axis. Gates used for selection are displayed as black outline boxes. Selection criteria for microglia ($CD11b^{+}$ $CD45^{lo}$) and macrophage ($F4/80^{+}$) are as described in the Flow Cytometry section in Methods. (a) Unstained brain cell suspension. Minimal recorded events within the gate can be attributed to background fluorescence; (b) Stained brain cell suspension with microglia identified in the gated area in purple; (c) Unstained alveolar lavage cell suspension. Again minimal recorded events within the gated area attributable to background; (d) Stained alveolar lavage cell suspension with macrophages identified in the gate in red.
