## Supplementary figures and images for "Defining the pig microglial transcriptome reveals their core signature, regional heterogeneity, and similarity with humans"

### Figure S2

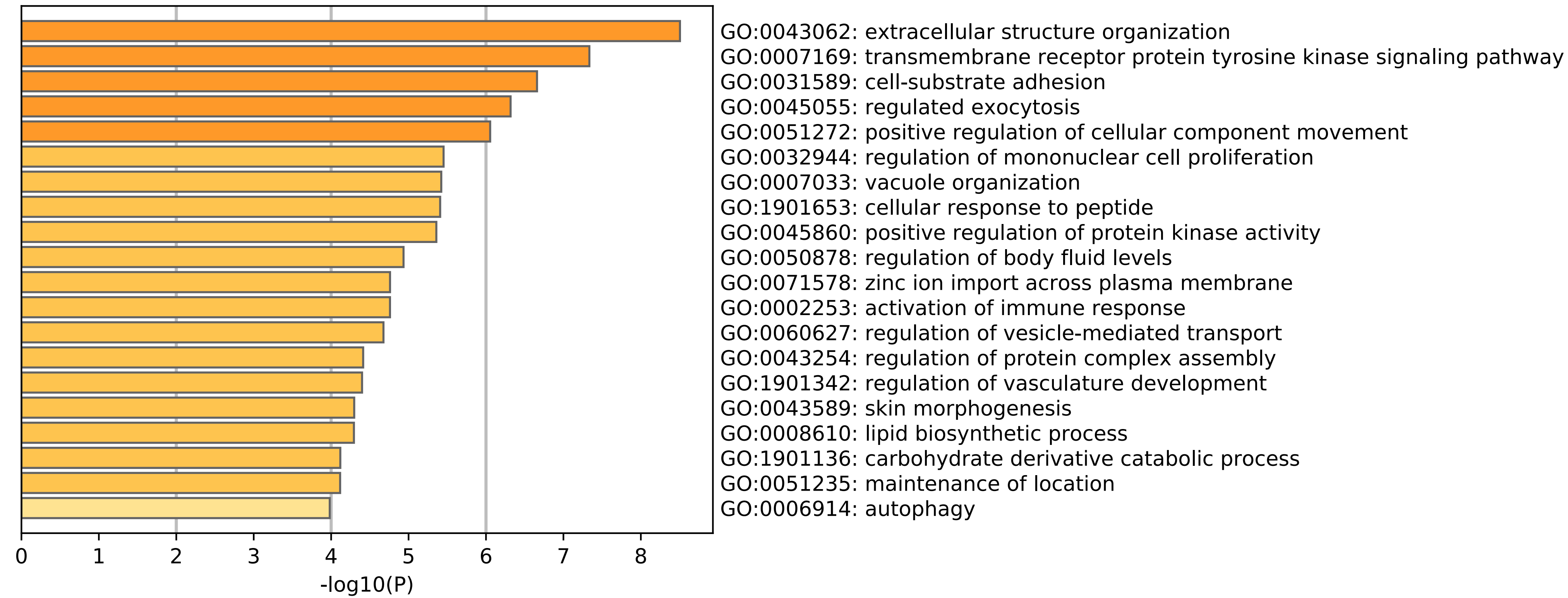

### Figure S3

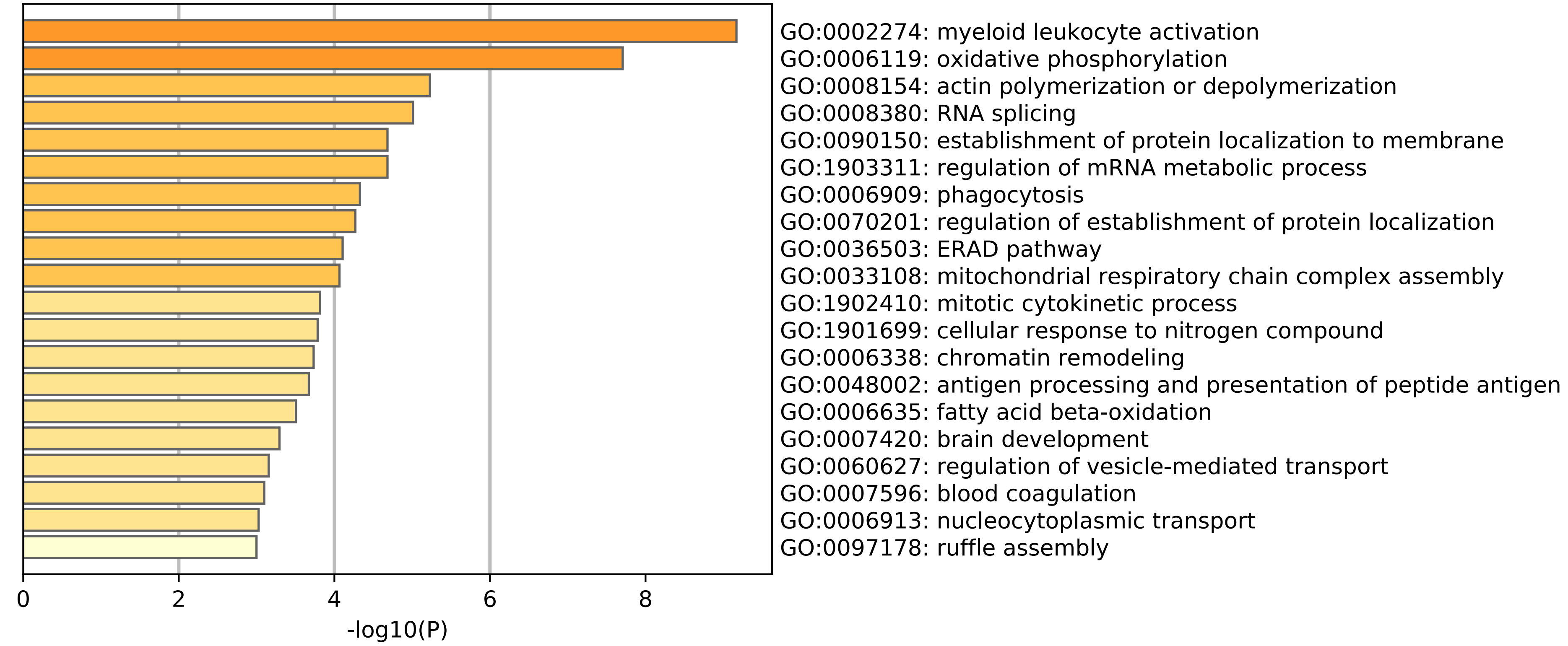

### Figure S4

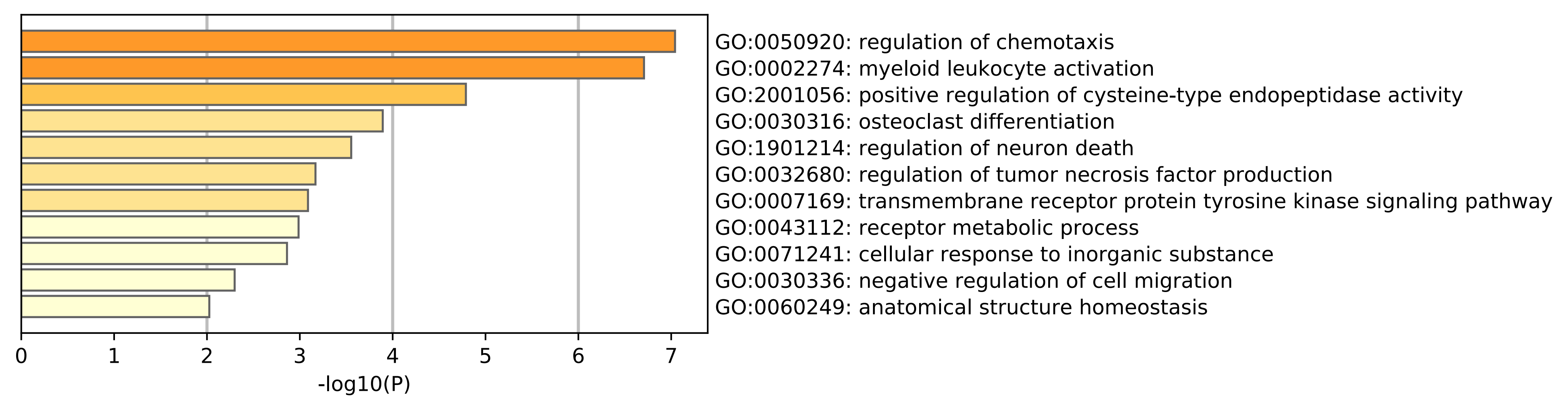

### Figure S5

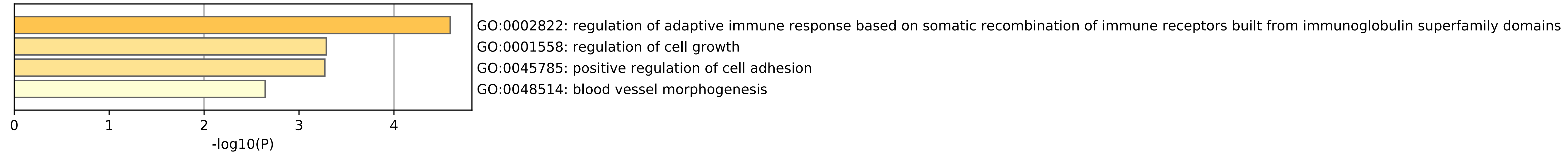
